# Type I interferons act directly on nociceptors to produce pain sensitization: Implications for viral infection-induced pain

**DOI:** 10.1101/2019.12.27.889568

**Authors:** Paulino Barragan-Iglesias, Úrzula Franco-Enzástiga, Vivekanand Jeevakumar, Andi Wangzhou, Vinicio Granados-Soto, Zachary T. Campbell, Gregory Dussor, Theodore J. Price

## Abstract

One of the first signs of viral infection is body-wide aches and pain. While this type of pain usually subsides, at the extreme, viral infections can induce painful neuropathies that can last for decades. Neither of these types of pain sensitization are well understood. A key part of the response to viral infection is production of interferons (IFNs), which then activate their specific receptors (IFNRs) resulting in downstream activation of cellular signaling and a variety of physiological responses. We sought to understand how type I IFNs (IFN-α and IFN-β) might act directly on nociceptors in the dorsal root ganglion (DRG) to cause pain sensitization. We demonstrate that type I IFNRs are expressed in small/medium DRG neurons and that their activation produces neuronal hyper-excitability and mechanical pain in mice. Type I IFNs stimulate JAK/STAT signaling in DRG neurons but this does not apparently result in PKR-eIF2α activation that normally induces an anti-viral response by limiting mRNA translation. Rather, type I interferons stimulate MNK-mediated eIF4E phosphorylation in DRG neurons to promote pain hypersensitivity. Endogenous release of type I IFNs with the double stranded RNA mimetic poly(I:C) likewise produces pain hypersensitivity that is blunted in mice lacking MNK-eIF4E signaling. Our findings reveal mechanisms through which type I IFNs cause nociceptor sensitization with implications for understanding how viral infections promote pain and can lead to neuropathies.

**SIGNIFICANCE STATEMENT:** It is increasingly understood that pathogens interact with nociceptors to alert organisms to infection as well as to mount early host defenses. While specific mechanisms have been discovered for diverse bacteria and fungal pathogens, mechanisms engaged by viruses have remained elusive. Here we show that type 1 interferons, one of the first mediators produced by viral infection, act directly on nociceptors to produce pain sensitization. Type I interferons act via a specific signaling pathway (MNK-eIF4E signaling) that is known to produce nociceptor sensitization in inflammatory and neuropathic pain conditions. Our work reveals a mechanism through which viral infections cause heightened pain sensitivity

## INTRODUCTION

Among the earliest symptoms of viral infection are aches and pain, effects that are usually body-wide suggest a systemic factor as a cause. While viral infections often cause pain that persists during the course of the ensuing illness, some viral infections, and sustained antiviral responses, can cause neuropathies leading to chronic pain (Brizzi and Lyons, 2014; Rodriguez et al., 2019). For instance, HIV and herpes viruses cause painful neuropathies that can last for decades (Hadley et al., 2016; Aziz-Donnelly and Harrison, 2017). Surprisingly little is known about the mechanisms through which viruses can induce acute pain and/or lead to neuropathies. One potential mechanism is through the upregulation of indoleamine 2,3 deoxygenase (IDO1) and subsequent increase in kynurenine signaling. In support of this idea, mice without the IDO1 enzyme lack hyperalgesia responses to certain viral infections (Huang et al., 2016). Another possibility is that early defense responses to viral infection trigger pain hypersensitivity. From this perspective, an ideal candidate is type I interferons (IFNs) because these cytokines are rapidly induced in a wide variety of cells upon exposure to virus. These IFNs then act via their cognate receptors to induce signaling in target cells (Schreiber, 2017; Barrat et al., 2019). We hypothesized that type I IFNs might act directly on peripheral nociceptors to cause pain.

A key component of the endogenous antiviral response is induction of cellular signaling that protects cells from viral infection and prevents viral replication. This is largely mediated by gene expression regulation signaling by type I IFNs. Type I IFNs alter gene expression in target cells by binding to heterodimeric transmembrane receptors composed of IFN receptor (IFNAR) 1 and 2 subunits and then engaging downstream signaling that activates transcriptional and translational programs in target cells (Levy and Darnell, 2002; Schreiber, 2017). The canonical IFNAR signaling pathway involves activation of janus kinase (JAK) and signal transducer and activation of transcription (STAT) -mediated changes in transcription (Levy and Darnell, 2002; de Weerd and Nguyen, 2012; Stark and Darnell, 2012). IFNAR activation also regulates translation of mRNAs through at least 3 pathways: 1) protein kinase R (PKR) driven phosphorylation of eukaryotic initiation factor 2 alpha (eIF2α) causing suppression of cap-dependent translation (Pindel and Sadler, 2011; Walsh et al., 2013), 2) phosphoinositide 3 kinase (PI3K) driven activation of the mechanistic target of rapamycin complex 1 (mTORC1) pathway augmenting translation of terminal oligopyrimidine tract (TOP) containing mRNAs (Thyrell et al., 2004; Hjortsberg et al., 2007), and 3) activation of extracellular signal regulated kinase (ERK) and mitogen activated protein kinase interacting kinase (MNK) signaling resulting in eIF4E phosphorylation and translation of mRNAs targeted by this phosphorylation event such as IFN stimulated genes (e.g. *Isg15* and *Isg54*), cytokines and matrix metalloproteinases (Platanias, 2005; Sen and Sarkar, 2007; Joshi et al., 2009). MNK-eIF4E activation is engaged by type I IFNs through the canonical IFNAR-JAK signaling pathway suggesting that STAT-mediated transcriptional changes and MNK-eIF4E-driven translation changes act in concert during the endogenous antiviral response (Joshi et al., 2009). Therefore, type I IFNs produce a direct antiviral effect through PKR-mediated eIF2α phosphorylation to suppress translation and block viral replication and an indirect effect via activation of MNK-eIF4E-mediated translation to augment host defense strategies such as increased immune surveillance (Joshi et al., 2009; Pindel and Sadler, 2011; Munir and Berg, 2013).

Nociceptors are tuned to detect a vast variety of immune modulators and can play a key role in host defense by responding directly to pathogenic organisms (Chiu et al., 2016; Foster et al., 2017; Pinho-Ribeiro et al., 2017). In response to pathogens or inflammatory mediators, nociceptors change their sensitivity generating nociceptive signals that act as a warning system (Liu et al., 2012; Baral et al., 2016). Overactivation of this system can lead to the generation of chronic pain disorders and damage to these neurons can cause neuropathic pain (Pinho-Ribeiro et al., 2017; Malcangio, 2019; Rodriguez et al., 2019). A key pathway linking initial nociceptor activation to nociceptor hypersensitivity and potentially the development of chronic pain is engagement of translation regulation signaling (Obara and Hunt, 2014; Khoutorsky and Price, 2018; de la Pena et al., 2019). Importantly, eIF2α phosphorylation, mTORC1 activation and MNK-eIF4E signaling can all lead to persistent sensitization of nociceptors and all of these pathways have been linked to neuropathic pain disorders (Inceoglu et al., 2015; Khoutorsky et al., 2016; Moy et al., 2017; Megat et al., 2019; Shiers et al., 2019)

We have tested the hypothesis that type I IFNs generate a pain response via a direct action on nociceptors. We find compelling evidence that exogenous and endogenous type I IFNs produce mechanical hypersensitivity via MNK-eIF4E signaling in nociceptors. This likely occurs as a downstream consequence of canonical JAK signaling. We find no evidence for engagement of PKR-eIF2α signaling in nociceptors by type I IFNs. Our findings provide a mechanistic link between type I IFNs and MNK-eIF4E signaling as a causative factor in pain produced by viral infection.

## MATERIALS AND METHODS

### Animals

Male eIF4E^S209A^ and MNK1^−/−^ mice were a gift of the Sonenberg laboratory at McGill University (Ueda et al., 2004; Furic et al., 2010) and bred at the University of Texas at Dallas (UTD) to generate experimental animals. Genotype was confirmed at weaning using DNA from ear clips. Experimental C57BL/6J wild-type (WT) animals were obtained from an internally maintained C57BL/6J colony at UTD. Electrophysiological experiments using WT mice were performed using mice between the ages of 4 and 6 weeks at the start of the experiment. Behavioral experiments using *eIF4E*^*S209A*^, MNK1^−/−^ (knockout for the *Mknk1* gene) and WT mice were performed using mice between the ages of 8 and 12 weeks, weighing ∼20–25 g. All animal procedures were approved by the Institutional Animal Care and Use Committee at The University of Texas at Dallas and were performed in accordance with the guidelines of the International Association for the Study of Pain.

## Antibodies and chemicals

Rabbit primary antibodies against p-eIF4E^S209^ (cat # ab76256, 1:500) and p-PKR (ab32036, 1:1000) were procured from Abcam. Mouse anti-NeuN antibody (cat # MAB377, 1:500) was obtained from Millipore. Chicken (for ICC, cat # CPCA-Peri, 1:500) and mouse (for IHC, cat # MAB1527, 1;1000) primary antibodies against peripherin were obtained from Encor Biotechnology Inc. and Sigma-Aldrich, respectively. Rabbit primary antibodies against p-JAK1 (cat # 3331S, 1:1000), JAK1 (cat # 3332S, 1;1000), p-STAT1 (cat # 9171, 1:000), STAT1 (cat # 9172S, 1:1000), p-STAT3 (cat # 9134S, 1:1000), STAT3 (cat # 91325, 1:1000), p-mTOR (cat # 2976S, 1:1000), mTOR (cat # 2983S, 1:1000), BiP (cat # 3177, 1:1000), PKR (cat # 3072S, 1:1000), p-eIF2α^Ser51^ (cat # 9721, 1:1000), eIF2α (cat # 9722, 1: 1000), p-ERK (cat # 9101S, 1:1000), ERK (cat # 9102S, 1:1000), p-AKT (cat # 2965S, 1:1000), AKT (cat # 4691S, 1:1000), p-RS6 (cat # 2317, 1:1000), RS6 (cat # 2317, 1:1000), eIF4E (cat # 9742S, 1:1000), β-actin (cat # 4967S; 1:10,000), and GAPDH (cat # 2118, 1:10,000) were obtained from Cell Signaling Technology. Alexa Fluor- and HRP-conjugated secondary antibodies were obtained from Life Technologies. The integrated stress response inhibitor, ISRIB (cat # SML0843), was purchased from Sigma. Recombinant mouse IFN-α protein (cat # 12100-1, lot # 6454) was procured from R&D Systems. Recombinant mouse IFN-β protein (cat # IF011, lot # SLBX5164) and poly (I:C) (cat # P1530) were purchased from Millipore-Sigma.

### Primary cell culture of mouse DRG neurons and Western blot

Cultured primary DRG neurons were used to test the effects of IFN-α [(300 units (U)/mL] and IFN-β [(300 units (U)/mL] application. For these experiments, mice were anesthetized with isoflurane and killed by decapitation. Then DRGs, from all spinal levels, were dissected and placed in chilled Hanks balanced-salt solution (HBSS, Invitrogen) until processed. Dorsal root ganglia were digested in 1 mg/mL collagenase A (Roche, Mannheim, Germany) for 25 minutes at 37°C then subsequently digested in a 1:1 mixture of 1 mg/mL collagenase D and papain (Roche) for 20 minutes at 37°C. After this step, dorsal root ganglia were then triturated in a 1:1 mixture of 1 mg/mL trypsin inhibitor (Roche) and bovine serum albumin (BioPharm Laboratories, Bluffdale, UT), then filtered through a 70 μm cell strainer (Corning, NY). Cells were pelleted then resuspended in DMEM/F12 with GlutaMAX (Thermo Fisher Scientific, MA) containing 10% fetal bovine serum (FBS; Thermo Fisher Scientific), 5 ng/mL NGF, 1% penicillin and streptomycin, and 3 mg/mL 5-fluorouridine with 7 mg/mL uridine to inhibit mitosis of non-neuronal cells and were distributed evenly in a 6-well plate coated with poly-D-lysine (Becton Dickinson, NJ). DRG neurons were maintained in a 37°C incubator containing 5% CO2 with a media change every other day. On day 6, DRG neurons were treated with either vehicle, IFN-α or IFN-β for 1, 3, 6 and 24h. Following treatments, cells were rinsed with chilled 1X PBS buffer, harvested in 200 μL of RIPA lysis buffer (50□JmM Tris, pH 7.4, 150□JmM NaCl, 1□JmM EDTA, pH 8.0, and 1% Triton X-100) containing protease and phosphatase inhibitors (Sigma-Aldrich), and then sonicated for 5 seconds. To clear debris, samples were centrifuged at 14,000 rpm for 15 min at 4°C. Ten to 15 μg of protein was loaded into each well and separated by a 10% SDS-PAGE gel. Proteins were transferred to a 0.45 PVDF membrane (Millipore, MA) at 30 V overnight at 4°C. Subsequently, membranes were blocked with 5% non-fat dry milk in 1X Tris buffer solution containing Tween 20 (TTBS) for at least 2 h. Membranes were washed 3 times for 10 minutes each (3 x 10) in 1X TTBS, then incubated with primary antibodies overnight at 4°C. The following day, membranes were washed 3 x 10 each then incubated with the corresponding secondary antibodies at room temperature for 1 h. After incubation, membranes were washed with 1X TTBS 3 x 10 each. Signals were detected using Immobilon Western Chemiluminescent HRP Substrate (Millipore) and then visualized with Bio-Rad ChemiDoc Touch. Membranes were stripped using Restore Western Blot Stripping buffer (Thermo Fisher Scientific) and reprobed with another antibody. Analysis was performed using Image lab 6.0.1 software for Mac (Bio-Rad, Hercules, CA).

To perform western blot analysis (WB) from tissues, animals were anesthetized with isoflurane and killed by decapitation. Lumbar spinal dorsal horn, L4-L5 DRGs and sciatic nerve were immediately frozen on dry ice and then sonicated for at least 10 seconds in 300 μL of RIPA lysis buffer containing protease and phosphatase inhibitors. To clear debris, samples were centrifuged at 14,000 rpm for 15 min at 4°C. Samples were processed for WB using the experimental protocol and analysis described above.

### Immunofluorescence

For primary neuronal cultures, DRG neurons were harvested and cultured for 6 days according to the protocol described above with the exception that cells were distributed evenly on poly-D-lysine–coated 8-well chamber slide with removable wells (cat # 12-565-8, Fisher Scientific). After treatments, cells were fixed in ice-cold 10% formalin in 1X PBS for 1 h and processed for immunocytochemistry (ICC). Cells were washed with 1X PBS and permeabilized in 1X PBS containing 10% heat-inactivated normal goat serum (NGS; Atlanta Biologicals, Atlanta, GA) and 0.02% Triton X-100 (Sigma) in 1X PBS for 30 min and then blocked in 10% NGS in 1X PBS for 2 h. Primary antibodies were applied overnight at 4°C and the next day appropriate secondary antibodies (Alexa Fluor; Invitrogen) were applied for 1 h. After additional 1X PBS washes, coverslips were mounted on Superfrost plus slides with ProLong Gold antifade (Invitrogen). Images were taken using an Olympus FluoView 1200 confocal microscope and analyzed with FIJI for Mac OS X. Images shown are representative of samples taken from 3 separate wells and presented as projections of Z stacks. Using ImageJ, the corrected total cell fluorescence (CTCF) was calculated to determine the intensity of the signal between experimental groups. To do so, the integrated density and the area, as well as the background noise was measured and the CTCF calculated as equal to: the integrated density - (area of selected cell x mean fluorescence of background readings). CTCF values from all experimental treatment groups were normalized to vehicle groups and expressed as normalized CTCF.

For tissues, animals were anesthetized with isoflurane and killed by decapitation, and tissues were frozen in O.C.T. on dry ice. Spinal cords were pressure ejected using chilled 1X PBS. Sections of L4-L5 DRGs (20 μm) were mounted onto SuperFrost Plus slides (Thermo Fisher Scientific) and fixed in ice-cold 10% formalin in 1X PBS for 1 h then subsequently washed 3 x 10 each in 1X PBS and processed for immunohistochemistry (IHC). Slides were permeabilized in 50% ethanol for 30 min. After 30 min, slides were washed 3 x 10 each in 1X PBS. Tissues were blocked for at least 2 h in 1X PBS and 10% NGS. Primary antibodies against NeuN, peripherin, p-eIF4E^S209^ and eIF4E were applied and incubated with DRG sections on slides at 4°C overnight. Immunoreactivity was visualized after 1 h incubation with Alexa Fluor secondary antibodies at room temperature. Images were taken using an Olympus FluoView 1200 confocal microscope. Images are presented as projections of Z stacks, and they are representative of samples taken from 3 animals.

### Single cell data

Single cell mouse DRG sequencing data from previously published work (Li et al., 2016) was used to generate Figures 2A–2D. Seurat package 2.2.1 (Butler et al., 2018) was used to cluster the single-cell data and visualization (van der Maaten and Hinton, 2008).

### Patch-clamp electrophysiology

Cell cultures for patch clamp electrophysiology were prepared as previously described (Moy et al., 2017). Male C57BL/6J mice (average age of 43 days) were anesthetized with 5% isoflurane and sacrificed by decapitation. DRGs were dissected and placed in ice-cold HBSS (divalent free), and incubated at 37°C for 15 min in 20 U/mL Papain (Worthington, Lakewood, NJ) followed by 15 min in 3 mg/ml Collagenase Type II (Worthington). After trituration through a fire-polished Pasteur pipette of progressively smaller opening sizes, cells were plated on poly-D-lysine and laminin (Sigma, St. Louis, MO)—coated plates. Cells were allowed to adhere for several hours at room temperature in a humidified chamber and then nourished with Liebovitz L-15 medium (Life Technologies, Grand Island, NY) supplemented with 10% fetal bovine serum (FBS), 10 mM glucose, 10 mM HEPES and 50 U/ml penicillin/streptomycin. The following day (within 24 h of dissociation), changes in neuronal excitability were tested after incubating the neurons with IFN-α (300 U/mL) for 1 h. To do so, whole-cell patch-clamp experiments were performed using a MultiClamp 700B (Molecular Devices) patch-clamp amplifier and PClamp 9 acquisition software (Molecular Devices) at room temperature. Recordings were sampled at 20 kHz and filtered at 3 kHz (Digidata 1550B, Molecular Devices). Pipettes (outer diameter, 1.5 mm; inner diameter, 1.1 mm, BF150-110-10, Sutter Instruments) were pulled using a PC-100 puller (Narishige) and heat polished to 3–5 MΩ resistance using a microforge (MF-83, Narishige). Series resistance was typically 7 MΩ and was compensated up to 60%. Data were analyzed using Clampfit 10 (Molecular Devices). All neurons included in the analysis had a resting membrane potential more negative than −40 mV. The RMP was recorded 1–3 min after achieving whole-cell configuration. In current-clamp mode, cells were held at −60 mV and action potentials were elicited by injecting slow ramp currents from 100 to 700 pA with Δ 200 pA over 1 s to mimic slow depolarization. Only cells that responded to the ramp depolarization – at least one spike at the maximum 700 pA, were considered for further analysis. The pipette solution contained the following (in mM): 120 K-gluconate, 6 KCl, 4 ATP-Mg, 0.3 GTP-Na, 0.1 EGTA, 10 HEPES and 10 phosphocreatine, pH 7.4 (adjusted with N-methyl glucamine), and osmolarity was ~285 mOsm. The external solution contained the following (in mM): 135 NaCl, 2 CaCl2, 1 MgCl2, 5 KCl, 10 glucose, and 10 HEPES, pH 7.4 (adjusted with N-methyl glucamine), and osmolarity was adjusted to ~315 mOsm with sucrose.

### Behavior

Mice were housed on 12-hour light/dark cycles with food and water available *ad libitum*. Mice were randomized to groups from multiple cages to avoid using mice from experimental groups that were cohabitating. Sample size was estimated by performing a power calculation using G*Power (version 3.1.9.2). With 80% power and an expectation of *d* = 2.2 effect size in behavioral experiments, and α set to 0.05, the sample size required was calculated as n = 6 per group. We therefore sought to have at least n = 6 sample in all behavioral experiments. SD (set at 0.3) for the power calculation was based on previously published mechanical threshold data from our lab (Moy et al., 2017). Animals were habituated for 1 hour to clear acrylic behavioral chambers before beginning the experiment. For intraplantar injections, drugs were injected in a total volume of 25 μL through a 30-gauge needle. For intraperitoneal injections, drugs were administered in a volume of 100 μL Mechanical paw withdrawal thresholds in mice were measured using the up–down method (Chaplan et al., 1994) with calibrated von Frey filaments (Stoelting Company, WoodDale, IL). Thermal latency was measured using a Hargreaves device (IITC Life Science) (Hargreaves et al., 1988) with heated glass settings of 29°C, 40% active laser power, and 20 sec cutoff were used. The experimenter was blinded to the genotype of the mice and the drug condition in all experiments.

### Quantification and statistical analysis

All results are presented as the mean ± SEM. Statistical differences between 2 groups were determined by the Student *t* test. One-or 2-way analysis of variance, followed by Dunnett or Bonferroni test, was used to compare differences between more than 2 groups. Post-hoc testing for electrophysiology data used Fisher’s LSD test. Differences were considered to reach statistical significance when *P* < 0.05. Complete statistical analysis is detailed on table 1. The *N* for each individual experiment is described in the figure legends. Data analysis was performed using GraphPad Prism 8.0 (GraphPad Software).

**Table 1.**
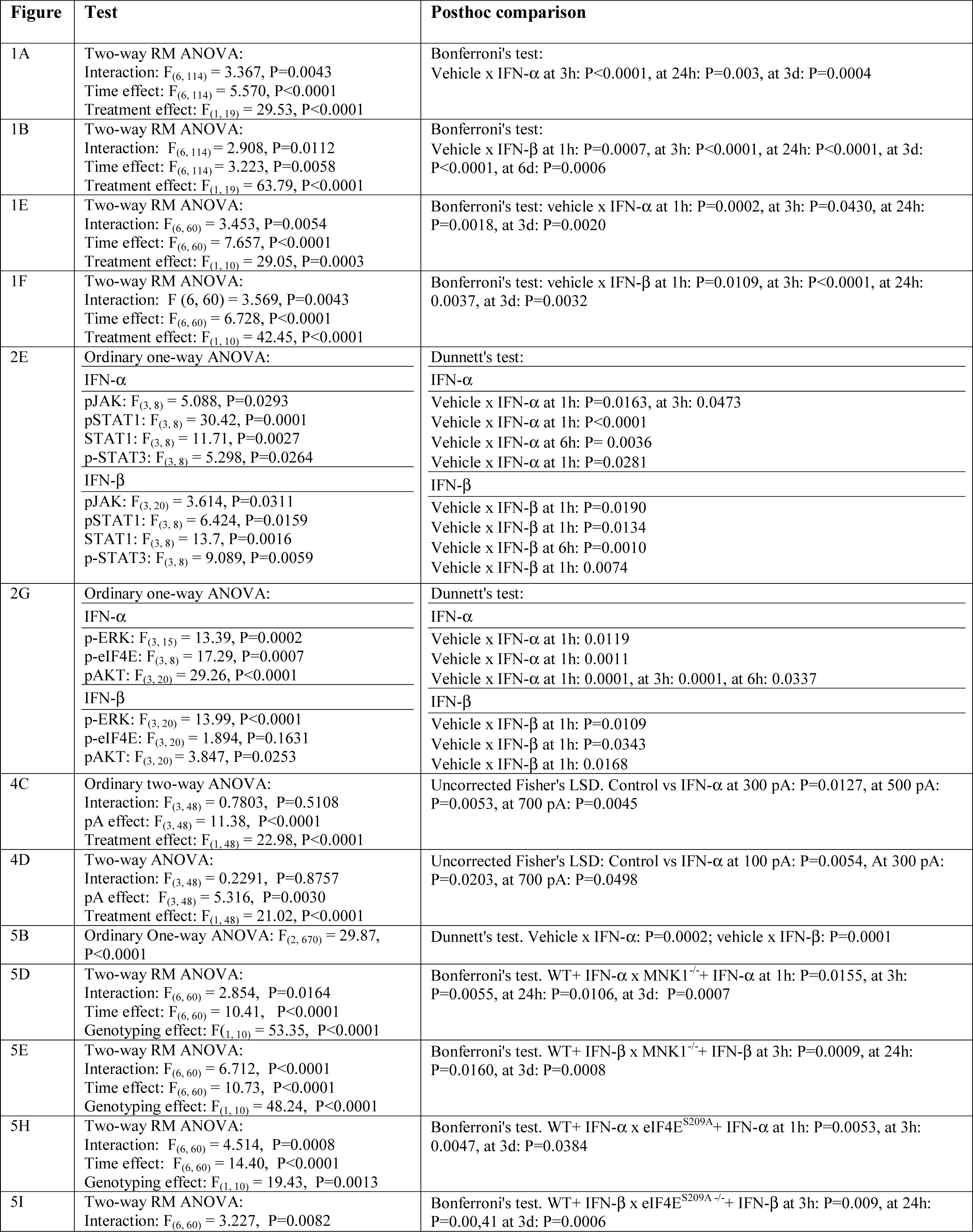
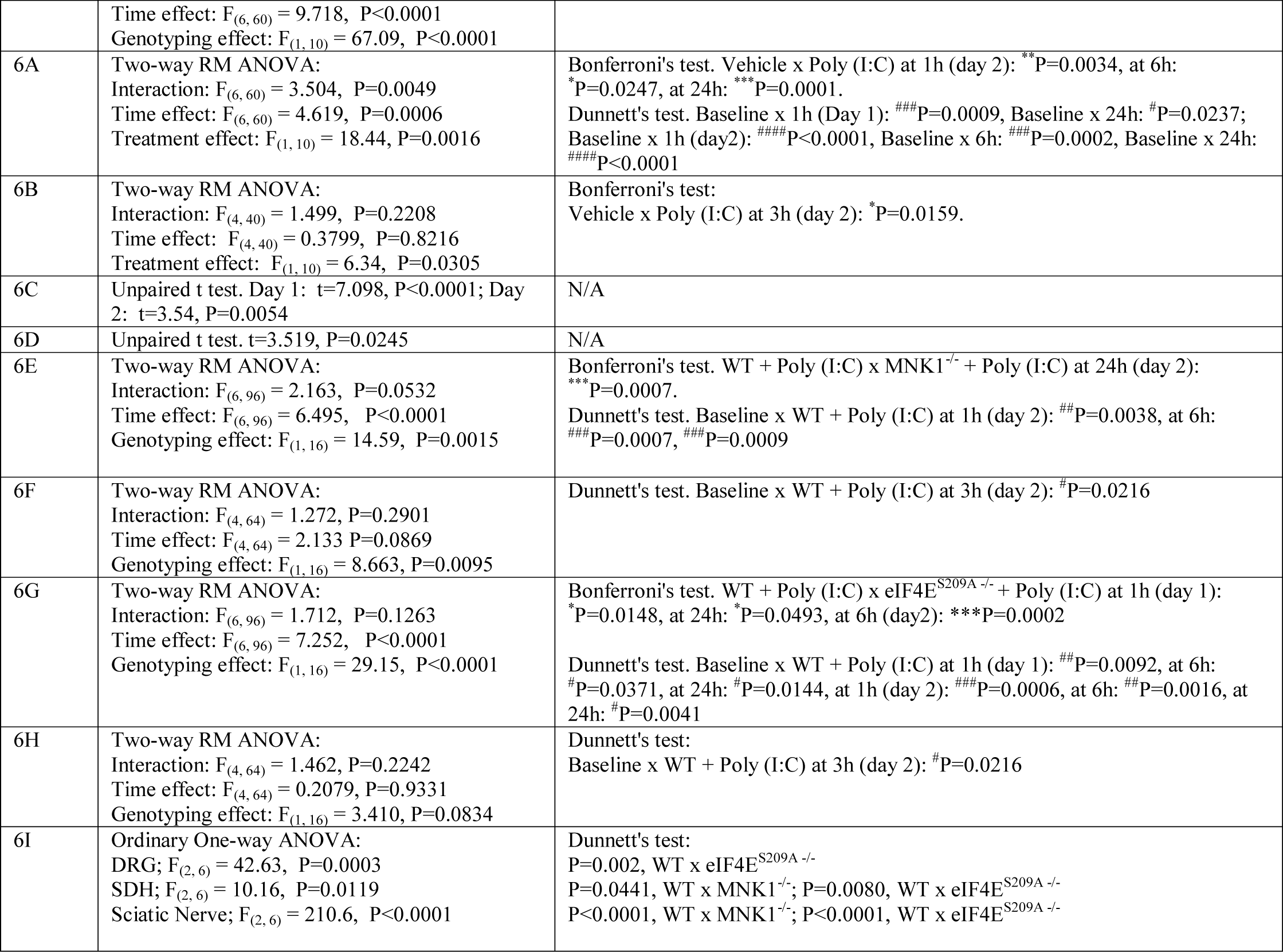
Student’s t-test and one- or two-way ANOVA with their respective *post-hoc* comparisons for each figure panel.

### Data and code availability

The data that support the findings of this study, including specific details of how tSNE plots were generated, are available from the corresponding author upon reasonable request.

## RESULTS

### Characterizing pain behavior responses induced by peripheral administration of type I IFNs

We first sought to investigate the nociceptive responses produced by type I interferons (α and β) *in vivo* in both sexes. The dose (300 units (U) – approximately 5 ng) of IFNs was chosen based on previous studies showing concentration-dependent effects on cellular signaling pathways (Larner et al., 1986; Hilkens et al., 2003) and studies showing plasma levels of type I IFNs in mice in response to viral infection (~ 1-2 ng/ml) (Gerlach et al., 2006; Shibamiya et al., 2009; Murray et al., 2015; Cheng et al., 2017). In male mice, intraplantar (i.pl.) administration of either IFN-α (300 U/25 μL) or IFN-β (300 U/25 μL), but not vehicle (saline), produced a rapid mechanical hypersensitivity, lasting for at least 3 days, to von Frey filament stimulation (Figures 1A and 1B) with no significant changes in paw withdrawal latency to thermal stimulation (Figures 1C and 1D). Likewise, in female mice, i.pl. administration of either IFN-α (300 U/25 μL) or IFN-β (300 U/25 μL) also increased hind paw mechanical hypersensitivity with no significant changes in thermal hypersensitivity (Figures 1G and 1H). No sex differences in the development of mechanical hypersensitivity (Figures 1I and 1J) or the presence of thermal hypersensitivity (Figures 1K and 1L) were observed between male versus female mice following either IFN-α (300 U/25 μL) or IFN-β (300 U/25 μL) i.pl. administration. These results show that direct activation of IFNRs by IFN-α or IFN-β creates a pronociceptive state that is likely produced via a peripheral site of action.

**Figure 1.**
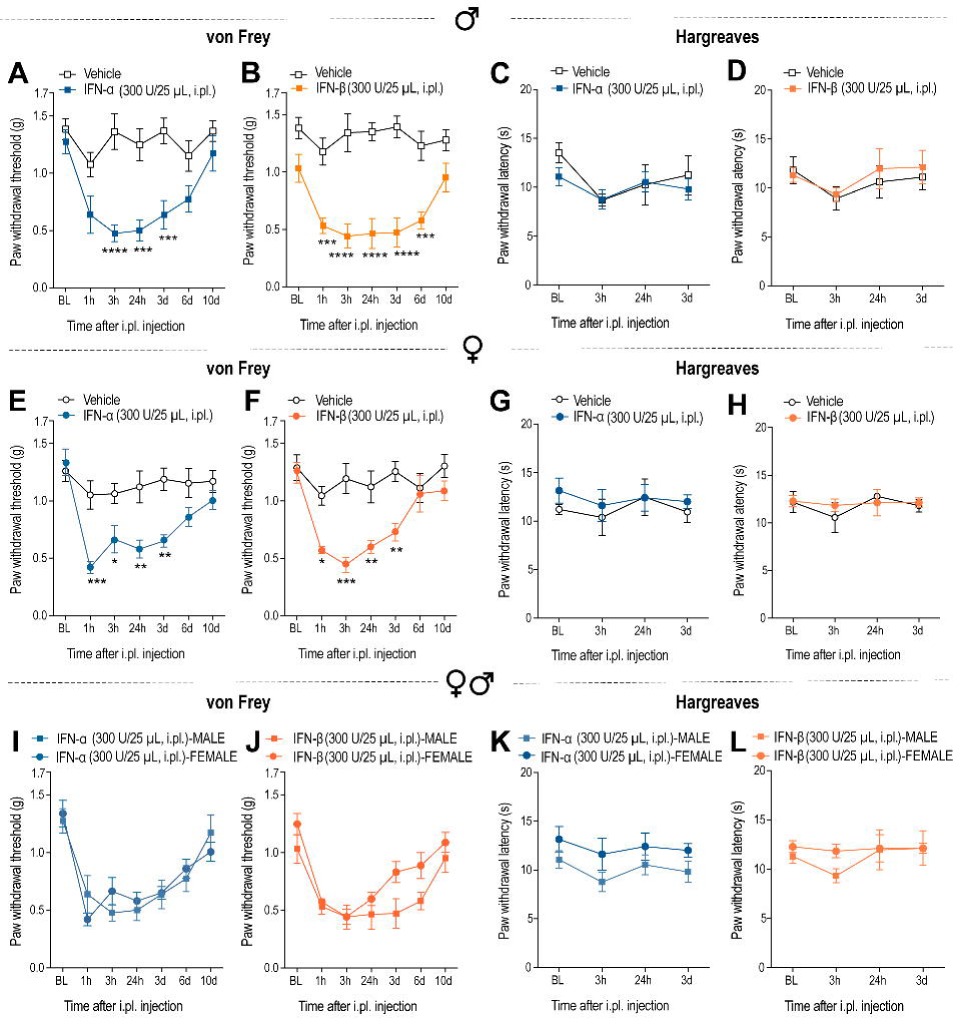
Type I IFNs induce mechanical nociceptive hypersensitivity responses via a peripheral action in male and female mice. (A-D) In male mice, intraplantar (i.pl.) administration of IFN-α (300 U/25 μL) or IFN-β (300 U/25 μL) increased paw mechanical hypersensitivity (g) to von Frey stimulation (A-B) with no significant changes in paw withdrawal latency (s) to thermal stimulation (C-D). n=9 (vehicle groups) and n=12 (IFN groups) in A-B. n=6 (vehicle groups) and n=12 (IFN groups) in C-D. (E-H) In female mice, i.pl. administration of IFN-α (300 U/25 μL) and IFN-β (300 U/25 μL) increased paw mechanical sensitivity to von Frey stimulation (E-F) with no significant changes in paw withdrawal latency (s) to thermal stimulation (G-H). n=6 per group. (I-L) When comparing male versus female, no significant sex differences in the development of mechanical hypersensitivity (I-J) or the presence of thermal sensitivity (K-L) were observed between groups following either IFN-α (300 U/25 μL) or IFN-β (300 U/25 μL) i.pl. administration. Data are presented as mean±SEM. Group differences were assessed using two-way ANOVA for A-B and E-F followed by Bonferroni’s multiple comparisons test. *p < 0.05, **p < 0.01, ***p < 0.001, ****p < 0.001.

### Identifying IFNRs expression in sensory neurons and their downstream signaling pathways

Because we observed a pronociceptive effect of IFN-α and IFN-β, we investigated the expression of IFNRs in DRG neurons. We used mouse DRG deeply RNA sequenced single cell data generated by (Li et al., 2016). We generated tSNE plots to show genes expression in specific clusters of cells (van der Maaten and Hinton, 2008). IFN-α and IFN-β bind a heterodimeric transmembrane receptor termed the IFN-α receptor (IFNAR), which is composed of IFNAR1 and IFNAR2 subunits (Schreiber, 2017). We observed expression of both *Ifnar1* (Interferon receptor 1, IFNR1) and *Ifnar2* (Interferon receptor 2, IFNR2) mRNAs with the neuronal marker *rbfox3* (NeuN), indicating their presence in neuronal populations in the mouse DRG (Figure 2A). *Ifnar2* mRNA appears to be more highly expressed than *Ifnar1* mRNA in many cells. *Ifnar1* and *Ifnar2* mRNAs were co-expressed among neurons that are likely to be nociceptors because they also express *Prph* (peripherin) and *Scn10a* (Nav1.8) (Figure 2B). Additionally, we detected that *Ifnar1* and *Ifnar2* mRNAs are widely distributed across nociceptors of peptidergic [*Trpv1* (TRPV1), *Calca* (CGRP)] and non-peptidergic [*P2rx3* (P2×3)] nature (Figure 2C). *Ifnar2* mRNA shows higher expression levels than *Ifnar1* in neurons containing *F2rl1* (PAR2) and *Nppb* (NPPB) mRNAs (Figure 2D). Therefore, IFNRs are present in DRG neurons and their activation could modulate nociceptive signaling events.

**Figure 2.**
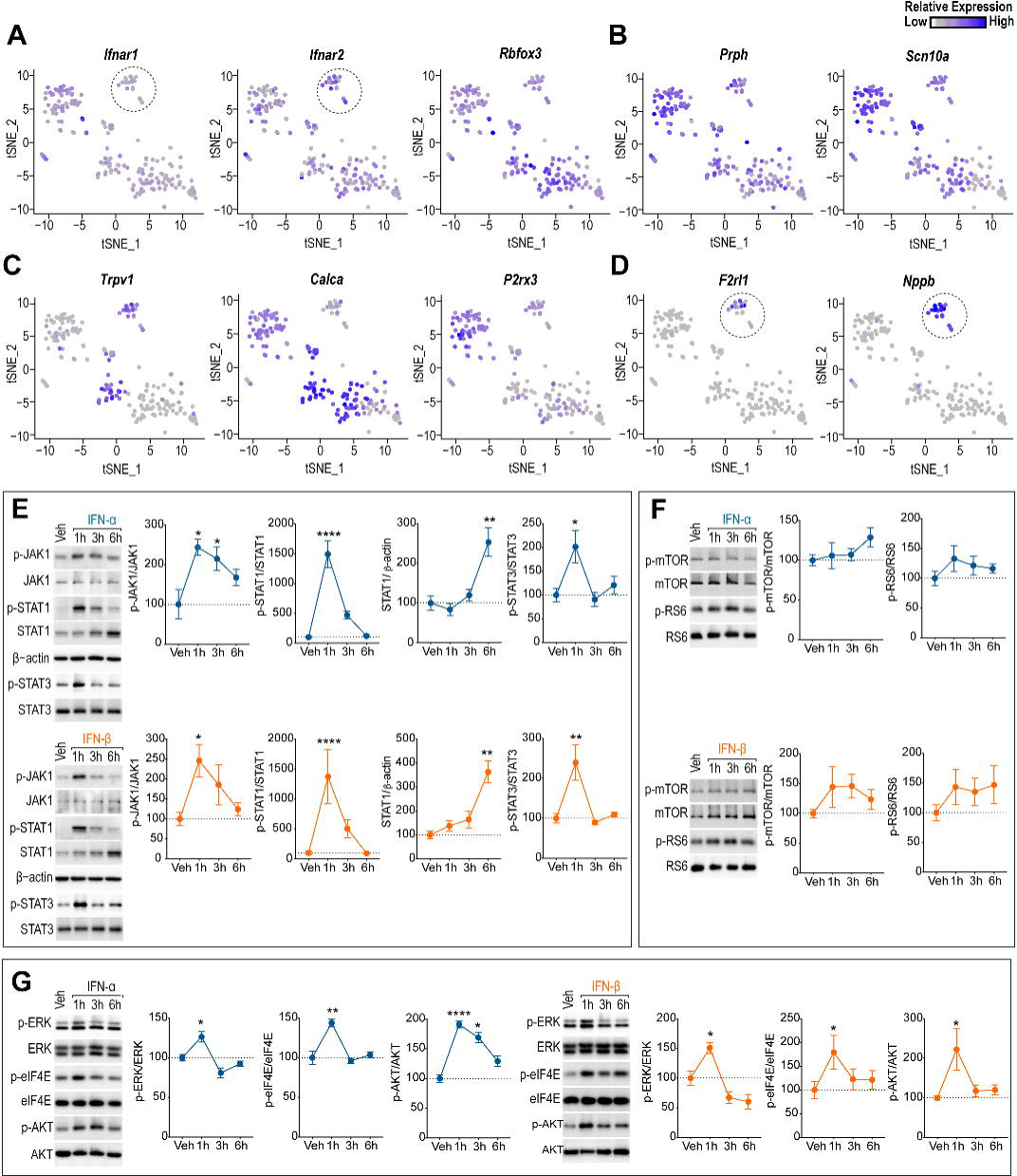
Expression of IFNRs in DRG sensory neurons and downstream signaling events associated with their activation. (A) Single DRG neuron-sequencing tSNE clusters showing expression of both *Ifnar1* (IFNR1) and *Ifnar2* (IFNR1) mRNAs with the neuronal marker *rbfox3* (NeuN). (B) *Ifnar1* and *Ifnar2* mRNAs expression overlaps with the small/medium-sized neuron sub-population expressing *Prph* (peripherin) and *Scn10a* (Nav1.8). (C) *Ifnar1* and *Ifnar2* mRNAs distribution across DRG sensory neurons of peptidergic [*Trpv1* (TRPV1), *Calca* (CGRP)] and non-peptidergic [P2×3 (P2×3)] sub-clusters. (D) *Ifnar1* and *Ifnar2* mRNAs expression differences in a subpopulation that express *F2rl1* (PAR2) and *Nppb* (NPPB), as indicated by stitched circles in A and D. (E) Direct stimulation of IFNRs with IFN-α (300 U/mL) and IFN-β (300 U/mL) activated downstream JAK/STAT signaling pathways in cultured DRG neurons. (F) Neither IFN-α (300 U/mL) nor IFN-β (300 U/mL) induced mTOR or RS6 phosphorylation in DRG cultures over a time course of 6 h. (G) Time course of the effects produced by IFN-α (300 U/mL) and IFN-β (300 U/mL) on ERK, eIF4E and AKT phosphorylation. Data are presented as mean±SEM. n=3-6 per group for WB analysis. Group differences (treated vs vehicle) in E-G were assessed using one-way ANOVA followed by Dunnett’s multiple comparisons test. *p < 0.05, **p < 0.01, ***p < 0.001, ****p < 0.0001

We then sought to investigate the downstream signaling events evoked by type I IFN application to cultured DRG neurons. We focused on two major pathways involved in type I IFN signaling in different cell types: transcriptional control via JAK/STAT (Levy and Darnell, 2002; de Weerd and Nguyen, 2012; Stark and Darnell, 2012) and translational control via two distinct pathways. Translation regulation by type I IFNs can occur through stimulation of cap-dependent translation via ERK/MAP kinase-MNK-eIF4E signaling (Walsh et al., 2013; Ivashkiv and Donlin, 2014) or inhibition of cap-dependent translation through induction of PKR-eIF2α signaling (Pindel and Sadler, 2011; Walsh et al., 2013) resulting in activation of the integrated stress response (ISR). Direct application of either IFN-α (300 U/mL) or IFN-β (300 U/mL) rapidly activated downstream JAK/STAT signaling pathways in cultured DRG neurons (Figure 2E). This signaling cascade involved the phosphorylation of JAK1, STAT1 and STAT3 together with a delayed increase in STAT1 total protein, likely representing a transcriptional change. In addition to STATs, other signaling factors have a role in IFN-mediated activities. These include activation of the AKT/mTOR/ULK1 pathway via PI3K and the ERK/MAP kinase pathway (Thyrell et al., 2004; Platanias, 2005; Hjortsberg et al., 2007; Saleiro et al., 2015). We did not observe any changes in mTOR or ribosomal protein S6 phosphorylation (Figure 2F) but we did observe an increase in ERK and eIF4E phosphorylation that occurred rapidly after type 1 IFN exposure (Figure 2G). Both IFN-α and IFN-β also stimulated AKT phosphorylation (Figure 2G). These findings demonstrate that type I IFNs engage cap-dependent translation regulation signaling via MNK-eIF4E.

Type 1 IFNs are also known to regulate translation via induction of PKR and activation of the integrated stress response (ISR). We did not observe changes in p-PKR or p-eIF2α levels and no changes were observed in BiP expression (ER chaperone protein) in response to type I IFN exposure (Figure 3A and 3B). Twenty four hr exposure to either IFN-α or IFN-β did not modify PKR phosphorylation or expression suggesting that type 1 IFNs do not induce PKR expression in DRG neurons (Figure 3C). In further support of these observations, no changes on p-PKR/PKR after a long IFN-α exposure were observed in the presence of the integrated stress response inhibitor ISRIB (Figure 3C), and ISRIB does not suppress signaling pathways shown to be modulated by IFN-α or IFN-β in our previous experiments (Figures 3D and 3E). Taken together, these experiments demonstrate that type I IFNs engage MNK-eIF4E signaling in DRG cultures and do not induce PKR activation to suppress cap-dependent translation via eIF2α phosphorylation.

**Figure 3.**
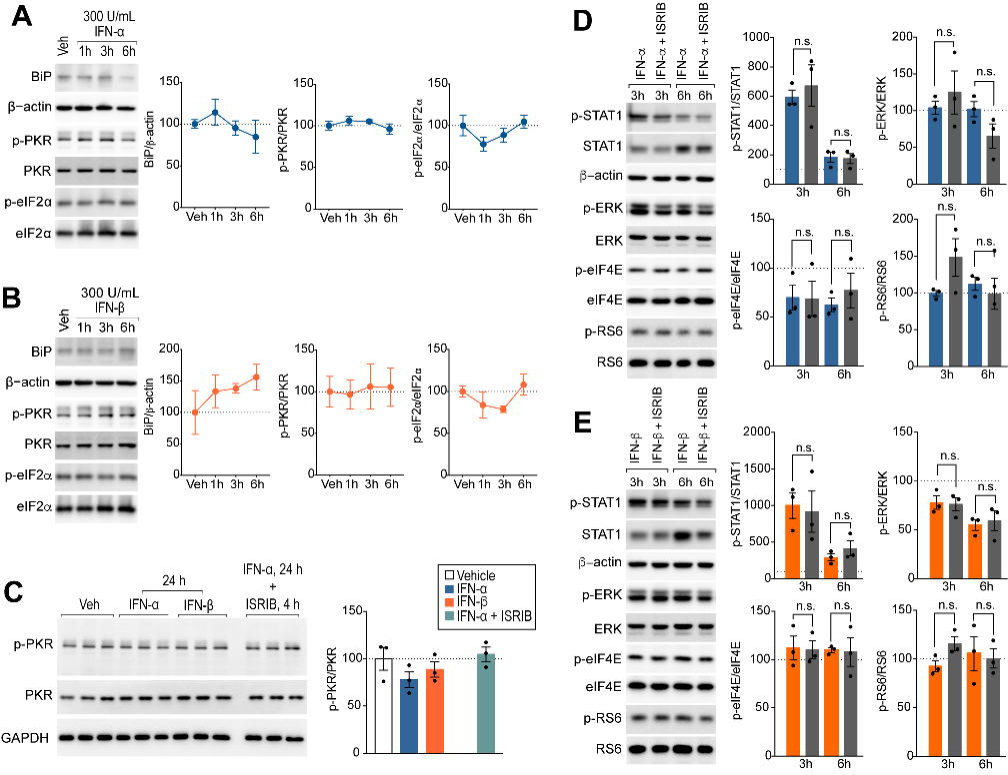
Type I IFNRs activity in cultured DRG neurons is neither associated with BiP-PKR-eIF2α stimulation nor suppressed by the small-molecule ISR inhibitor ISRIB. (A-B) Application of either IFN-α (300 U/mL) (A) or IFN-β (300 U/mL) (B) for 1-6 h did not modify Bip expression or signaling via PKR and downstream p-eIF2α in cultured DRG neurons. (C) Long exposure (24 h) to IFN-α (300 U/mL) (A) or IFN-β (300 U/mL) did not modify PKR phosphorylation in cultured DRG neurons. Likewise, no changes on PKR phosphorylation after a 24 h IFN-α (300 U/mL) treatment were observed in the presence of the integrated stress response inhibitor ISRIB (200 nM). (D-E) The integrated stress response (ISR) inhibitor ISRIB (200 nM) did not modulate components of the signaling pathways that are activated after either IFN-α (300 U/mL) (D) or IFN-β (300 U/mL) (E) application. n=3 per group. Data are presented as mean±SEM.

### Patch-clamp electrophysiology on DRG neurons links Type I IFNs activity to neuron hyperexcitability

To assess whether the effects of type I interferons contribute to nociceptor excitability, we exposed DRG neurons to IFN-α (300 U/mL) for ~1 h (average exposure time: 86.6±7 min) and measured neuronal excitability using patch-clamp electrophysiology. The treatment was present in both the L-15 culture medium and later in the external bath solution until completion of the electrophysiology experiments. Patch clamp electrophysiology was performed from small- and medium-sized populations of neurons in the cultured DRGs in both groups (capacitance: control 23.8 ± 2.9 pF vs IFN-α 24.3 ± 1.5 pF, P = 0.89; diameter: control 26.5 ± 0.56 pF vs IFN-α 26.8 ± 0.53 pF, P = 0.7; Figure 4A). Resting membrane potential (RMP) was more hyperpolarized than −40 mV in all cells sampled and IFN-α treatment did not alter the RMP compared to the control group (control −51.2 ± 3 mV vs IFN-α −47.5 ±2.5 mV, P = 0.38; Figure 4A). In response to ramp current injections mimicking slow depolarizations, DRG neurons exposed to IFN-α showed elevated excitability, measured as the number of action potentials elicited, compared to the control group with a significant main effect of treatment (F_(1,48)_ = 22.9, P < 0.001). Significant differences were observed at each time point of ramp injection tested (Figures 4B and 4C). We further measured the latency to the first spike following ramp current injection and determined that exposure to IFN-α shortened the latency of initiation of the action potential (F_(1,48)_ = 21.02, P < 0.001; Figure 4D). Therefore, type I IFN exposure rapidly promotes hyperexcitability in small diameter DRG neurons over a time course coinciding with MNK-eIF4E activation.

**Figure 4.**
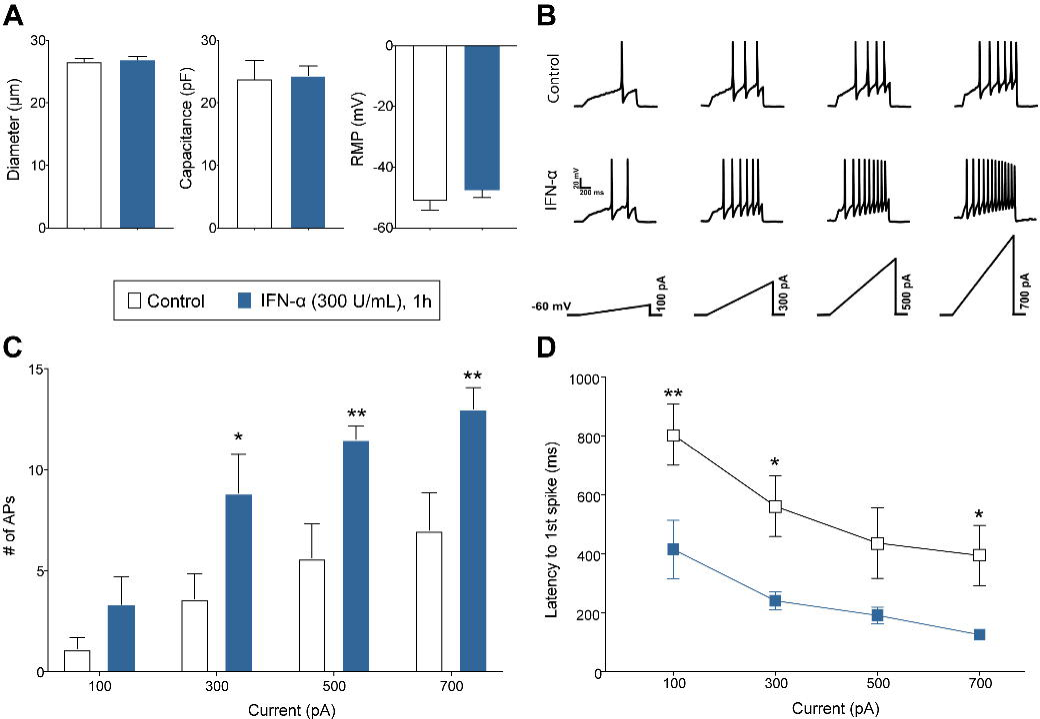
IFN-α treatment causes DRG neuron hyperexcitability. (A) Small- and medium sized DRG neurons were sampled for patch clamp electrophysiology experiments. The resting membrane potential was similar across the groups, with no significant effect of IFN-α treatment. (B) Representative traces of action potential firing in the control (n = 8 cells) and IFN-α (n = 6 cells) groups. Action potentials were elicited by slowly depolarizing ramp currents of varying intensities. (C) Mean number of action potentials were higher in the IFN-α group at each ramp intensity. (D) IFN-α treatment significantly shortened the latency to the first spike. Group differences were assessed using two-way ANOVA followed by Fisher’s LSD test. * P < 0.05, ** P < 0.01.

### MNK-eIF4E signaling links type I IFN actions on sensory neurons to mechanical hypersensitivity

Since we previously observed that ERK/MNK-eIF4E signaling axis was the primary component contributing to IFN-α and IFN-β effects in DRG neurons, we targeted this pathway using genetic tools to investigate its contribution to type 1 IFN-induced pain hypersensitivity. When ERK is activated, it subsequently phosphorylates MNK1/2 (Waskiewicz et al., 1997) leading to phosphorylation of eIF4E at serine 209 (Waskiewicz et al., 1999). We used immunocytochemistry (ICC) on cultured DRG neurons to assess whether type 1 IFNs impacts eIF4E phosphorylation at single cell resolution. We found that one-hour stimulation with either IFN-α (300 U/mL) or IFN-β (300 U/mL) stimulated phosphorylation of eIF4E mostly in neurons expressing peripherin, a marker for nociceptors (Figure 5A and 5B). To assess the behavioral impact of this signaling, we used MNK1^−/−^ mice (Figure 5C) and tested mechanical hypersensitivity after i.pl. IFN-α (300 U/25 μL) or IFN-β (300 U/25 μL) administration. Mechanical hypersensitivity was attenuated in MNK1^−/−^ mice compared to WT mice following IFN application (Figures 5D and 5E). Furthermore, mice lacking eIF4E phosphorylation at serine 209 (eIF4E^S209A^, Figure 5F) showed a complete absence of eIF4E phosphorylation in lumbar (L5) DRGs (Figure 5G) and a significant reduction in mechanical hypersensitivity following i.pl. IFN injection (Figures 5H and 5I). We conclude that a MNK-eIF4E signaling mechanism strongly contributes to type I IFN-induced pain hypersensitivity.

**Figure 5.**
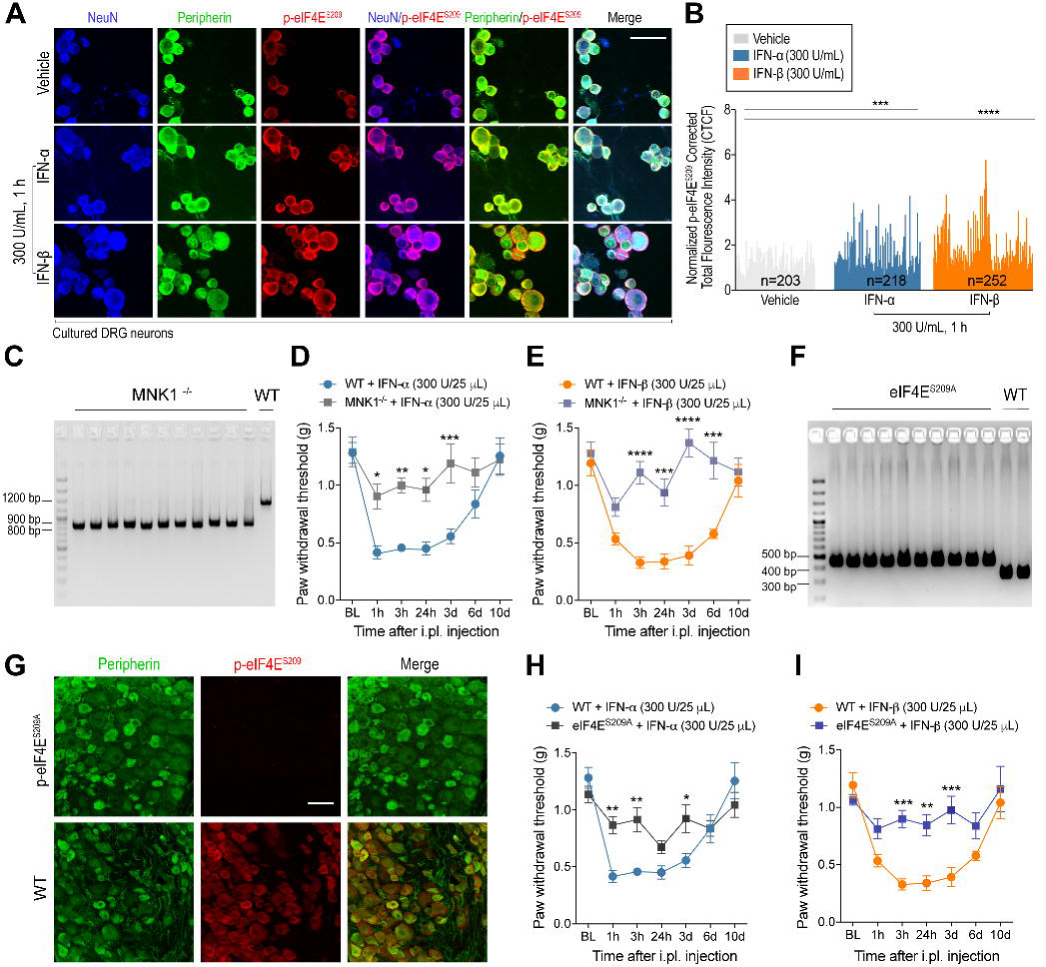
Genetic and pharmacological targeting of the MNK-eIF4E signaling axis attenuates type I IFN-induced pain hypersensitivity. (A-B) One h stimulation with either IFN-α (300 U/mL) or IFN-β (300 U/mL) phosphorylated the translation initiation factor eIF4E at serine S209 (eIF4E^S209^) in cultured DRG neurons expressing NeuN (blue) and peripherin (red). Scale bar 50 μm. Data are presented as mean±SEM. Group differences were assessed using one-way ANOVA followed by Dunnett’s multiple comparisons test. (C) MNK1^−/−^ mouse genotyping. (D-E) Mechanical hypersensitivity produced by intraplantar (i.pl.) administration of either IFN-α (300 U/25 μL) (D) or IFN-β (300 U/25 μL) (E) was attenuated in mice lacking MNK1 (*Mknk1*^−/−^), the specific kinase that phosphorylates eIF4E. Data are presented as mean±SEM. n=6 per group. Group differences were assessed using two-way ANOVA followed by Bonferroni’s multiple comparisons test. *p < 0.05, **p < 0.01, ***p < 0.001, ****p < 0.0001 (F) eIF4E^S209A^ mouse genotyping. (G) Mice lacking the phosphorylation site at Serine 209 (eIF4E^S209A^) showed absence of eIF4E phosphorylation in L5 DRGs demonstrating antibody specificity. Scale bar=50 μm. (H-I) Mechanical hypersensitivity was attenuated in eIF4E^S209A^ mice compared to WT mice following an i.pl. injection of either IFN-α (300 U/25 μL) (H) or IFN-β (300 U/25 μL) (I). Data are presented as mean±SEM. n=6 per group. Group differences were assessed using two-way ANOVA followed by Bonferroni’s multiple comparisons test. *p < 0.05, **p < 0.01, ***p < 0.001

### Induction of endogenous type I interferon response with poly (I:C) causes MNK-eIF4E-dependent pain hypersensitivity

Type I IFN responses are caused by viral infections and sustained elevations in type I IFNs have been associated with multiple autoimmune diseases including systemic lupus erythematosus (SLE) and rheumatoid arthritis (Forster, 2012). Moreover, therapeutic IFN-α administration has also been reported as associated with the emergence of somatic symptomatology such as body pain, myalgias, headache, joint pain, abdominal pain (Capuron et al., 2002; Shakoor et al., 2010; Nogueira et al., 2012), and inflammatory hyperalgesia (Fitzgibbon et al., 2019). To investigate how endogenous type I IFN production causes pain sensitization, we intraperitoneally (i.p.) injected mice, for 2 consecutive days, with a synthetic analog of a double stranded RNA (dsRNA), poly (I:C) (1 mg/kg). Poly (I:C) is well-known to activate a number of transcription factors, including IFN regulatory factor 3 (IRF3) resulting in the production of IFN-α and IFN-β (Kawai and Akira, 2008). We found that mice injected with poly (I:C) developed mechanical hypersensitivity (Figure 6A) as well as thermal hypersensitivity (Figure 6B) over a time-course of 3-24 h after the second poly (I:C) administration. Changes in mechanical and thermal hypersensitivity were preceded by an increase in core body temperature, consistent with known physiological effects of poly (I:C) (Figure 6C). Based on our previous observations, we hypothesized that the effects seen on thermal and mechanical hypersensitivity would be mechanistically linked to MNK-eIF4E signaling. As predicted, poly (I:C) administration increased phosphorylated eIF4E immunoreactivity in L5 DRGs of WT mice without affecting total eIF4E protein (Figure 6D). Mechanical (Figure 6E) and thermal (figure 6F) hypersensitivity produced by poly (I:C) was attenuated in MNK1^−/−^ compared to WT mice. Similarly, mechanical (Figure 6G) and thermal (Figure 6H) hypersensitivity was decreased in eIF4E^S209A^ mice compared to WTs. Moreover, L4-L5 DRGs, lumbar spinal dorsal horn (SDH) and sciatic nerve from MNK1^−/−^ and eIF4E^S209A^ mice showed a decrease and absence, respectively, of eIF4E phosphorylation compared to WT mice following poly (I:C) administration (Figure 6I). Finally, we tested whether poly (I:C) had a direct effect on DRG neurons. Direct application of poly (I:C) did not increase p-ERK, p-eIF4E, p-PKR or p-eIF2α in cultured DRG neurons (Figure 6J), suggesting that effects observed with poly (I:C) *in vivo* are unlikely explained by a direct action of the compound on DRG neurons. Instead, poly (I:C) likely acts via endogenous production of type I IFNs that then act on DRG neurons. These experiments demonstrate that endogenous type I IFN production acts via MNK-eIF4E signaling to induce pain hypersensitivity.

**Figure 6.**
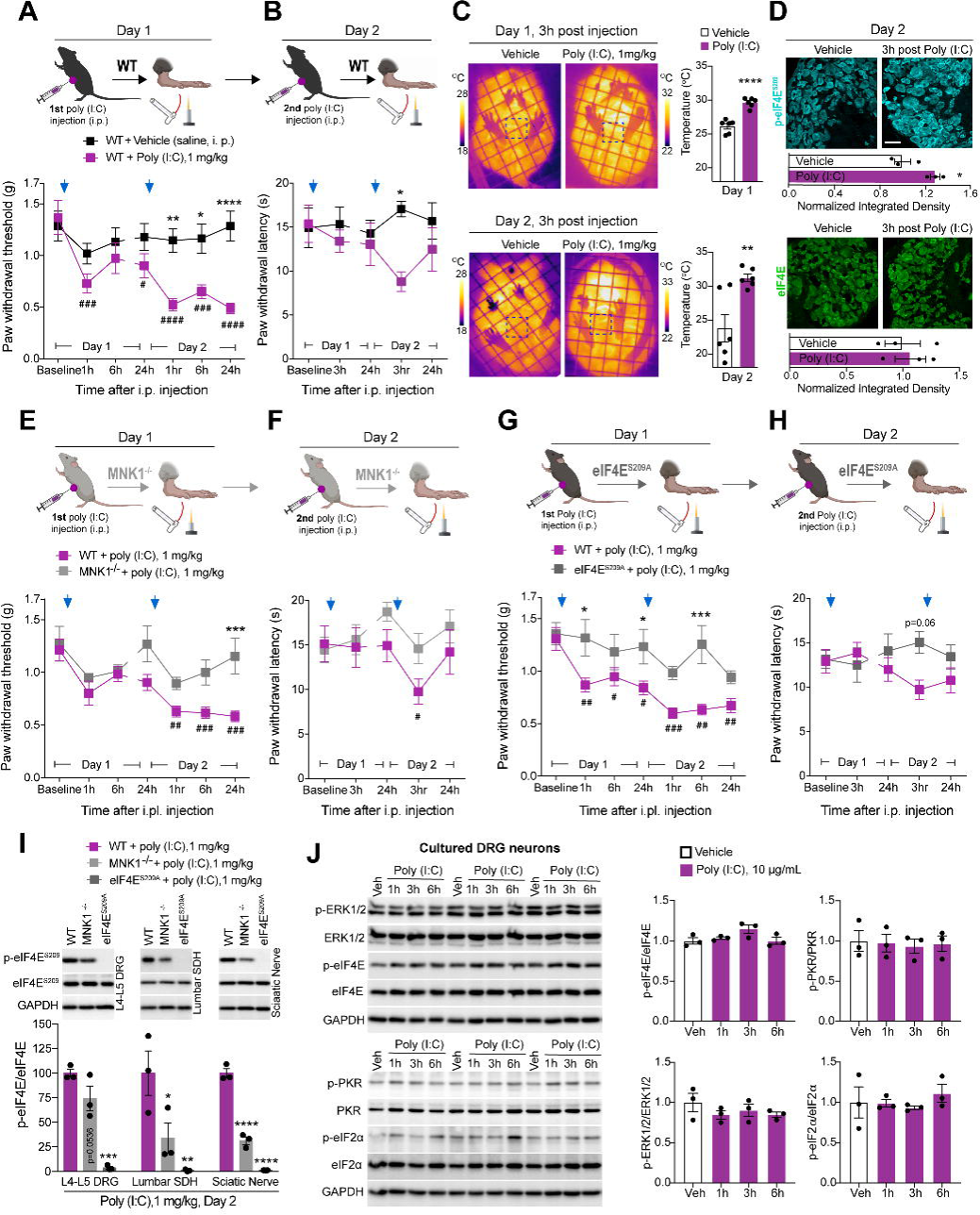
Endogenous type I IFN induction with poly (I:C) produces pain sensitization in mice via MNK1-eIF4E signaling. (A-B) The synthetic analog of a double stranded RNA (dsRNA), poly (I:C) (1mg/kg, i.p.), injected for two consecutive days, produced mechanical (A) and thermal hypersensitivity (B) in mice. n=6 per group. *p < 0.05, **p < 0.01, compare to vehicle. ^#^p < 0.05, ^##^p < 0.01, ^###^p < 0.001, ^####^p < 0.0001 compare to baseline. Group differences were assessed using two-way ANOVA followed by Dunnett’s (^#^) or Bonferroni’s (*) multiple comparisons tests. (C) Changes in body temperature produced by intraperitoneal poly (I:C) (1 mg/kg) administration. n=6 per group. **p < 0.01, ****p < 0.0001 compared to vehicle. Group differences were assessed using unpaired t-test. (D) Intraperitoneal poly (I:C) administration increased phospho, but not total, eIF4E in L5 DRGs of WT mice at day 2 (3 h post 2^nd^ poly I:C injection). Scale bar=50 μm. n=3 per group. *p < 0.05 compared to vehicle. Group differences were assessed using unpaired t-test. (E-F) Mechanical (E) and thermal (F) hypersensitivity produced by i.pl. administration of poly (I:C) (1 mg/kg) were partially attenuated in MNK1^−/−^ mice compared to WT mice. ***p < 0.001, compare to WT mice. ^#^p < 0.05, ^##^p < 0.01, ^###^p < 0.001, ^####^p < 0.001 compared to baseline. n=12 (WT) and n=6 (MNK1^−/−^) per group. Group differences were assessed using two-way ANOVA followed by Dunnett’s (^#^) or Bonferroni’s (*) multiple comparisons tests. (G-H) Mechanical (G) and thermal (H) hypersensitivity produced by i.pl. administration of poly (I:C) (1 mg/kg) were attenuated in eIF4E^S209A^ mice compared to WT mice. n=12 (WT) and n=6 (eIF4E^S209A^) per group. *p < 0.05, ***p < 0.001 compared to WT mice, ^#^p < 0.05, ^##^p < 0.01, ^###^p < 0.001 compared to baseline. Group differences were assessed using two-way ANOVA followed by Dunnett’s (^#^) or Bonferroni’s (*) multiple comparisons tests. (I) Lumbar DRGs (L4-L5), lumbar spinal dorsal horn (SDH) and sciatic nerve from MNK^−/−^ and eIF4E^S209A^ mice showed decrease and absence, respectively, on eIF4E phosphorylation compared to WT mice following i.p. poly I:C administration [day 2, 3 h post 2^nd^ poly (I:C) injection]. n=3 per group. *p < 0.05, **p < 0.01, ***p < 0.001, ****p < 0.001 compared to WT. Differences were assessed using one-way ANOVA followed by Dunnett’s multiple comparisons test. (J) Application of poly (I:C) (10 μg/mL) did not increase p-ERK, p-eIF4E, p-PKR or p-eIF2α in cultured DRG neurons. n= 3 per group.

## DISCUSSION

Our findings provide evidence for a mechanistic link between viral infection, type 1 IFN production and rapid induction of nociceptor hyperexcitability and mechanical pain sensitization. This occurs via a direct action of type I IFN receptors on sensory neurons and is dependent on downstream signaling via MNK-eIF4E. We find no evidence for mTORC1 activation or induction of eIF2α phosphorylation in DRG neurons by type I IFNs, demonstrating that the key translation regulation pathway engaged is eIF4E phosphorylation. Collectively, these results provide molecular insight into why one of the first signs of viral infection is body-wide aches and pain.

While it is well known that viral infection can cause pain, very little work has been done to understand the underlying mechanisms driving this effect (Chiu et al., 2016). Aches and pain caused by viral infection have classically been attributed to fever but these aches and pain often begin before the onset of fever. Our findings with poly (I:C) treatment in mice show that fever and pain effects are disassociated, but in this case the fever clearly preceded hyperalgesia caused by poly (I:C) treatment. An alternative mechanism for viral infection-induced pain is upregulation of IDO1 enzyme and consequent increased production of kynurenine. In support of this idea, mice lacking IDO1 show decreased pain sensitization in response to viral infection (Huang et al., 2016). However, subsequent work demonstrates that this IDO1 upregulation occurs via virally-mediated upregulation of IFN-β and that IDO1 expression can be driven by type I IFNs (Gaelings et al., 2017). Our work demonstrates that this initial type 1 IFN induction by viral infection can drive a direct sensitization of nociceptors through type 1 IFN receptors expressed by these sensory neurons. This suggests that early pain sensitization caused by viral infection may proceed independently of IDO1 upregulation. A direct effect of type I IFNs on nociceptors and type 1 IFN-mediated upregulation of IDO1 and subsequent kynurenine signaling may act in concert to cause prolonged pain responses that can occur with viral infections.

Our work adds to a growing understanding of how pathogens and host-defense responses interact with nociceptors (Chiu et al., 2016). Bacteria can act directly on nociceptors via N-formylated peptides that are agonists of G protein-coupled formyl peptide receptors (Chiu et al., 2013; Pinho-Ribeiro et al., 2017) that are expressed in mouse and human DRG neurons (Ray et al., 2018). Bacteria also release α-hemolysin which directly excites nociceptors to cause pain (Chiu et al., 2013; Blake et al., 2018). While nociceptors can detect bacterial invasion, rapidly sending an alert signal to the brain, they can also play more nuanced roles in bacterial host defense. For instance, gut-innervating nociceptors have very recently been shown to play an active role in defending against *Salmonella* infection. This happens via an effect of calcitonin gene-related peptide (CGRP) signaling on intestinal microvilli cells and resident microbiome to protect against *Salmonella* invasiveness (Lai et al., 2019). In the case of viruses, we find that a dsRNA mimetic, poly (I:C) does not seem to have a direct effect on nociceptors, suggesting that immune and other somatic cells are likely the first detectors of viral infection. Our findings clearly demonstrate, however, that one of the earliest responses to viral infection, production of type 1 IFN, causes robust and rapid sensitization of nociceptors via a specific translation regulation signaling cascade. Our findings regarding the action of type I IFNs on nociceptors are opposed to the recent work of Liu et al., (Liu et al., 2016) who found that IFNα causes inhibition of pain signaling at the level of the dorsal horn. These authors proposed that type I IFNs from astrocytes cause presynaptic inhibition of neurotransmitter release from nociceptors therefore reducing pain signaling. It is possible that CNS-released type I IFNs have a different action on nociceptor central terminals than type 1 IFNs released in the periphery have on nociceptor peripheral ending and cell bodies. Another possibility is that very high doses of type I IFNs (5,000-10,000 units) produce an inhibition of MAPK signaling, as recently shown in the dorsal horn in the context of neuropathic pain (Liu et al., 2019). We show that lower doses of type I IFNs (300 units) produce clear MAPK signaling activation in DRG neurons *in vitro* and *in vivo*. The dose used in our experiments is consistent with earlier studies examining dose-dependent effects of type I IFN signaling on what are now known as canonical signaling pathways (Larner et al., 1986; Hilkens et al., 2003) and with plasma levels induced by virus (Gerlach et al., 2006; Murray et al., 2015; Cheng et al., 2017) or poly (I:C) (Shibamiya et al., 2009).

Translation regulation is a central mechanism driving nociceptor hyperexcitability and mechanical pain (Khoutorsky and Price, 2018). A key antiviral response is activation of PKR and downstream phosphorylation of eIF2α (Balachandran and Barber, 2007). This results in decreased cap-dependent translation and suppression of viral replication capability in host cells. Another upstream eIF2α kinase, PERK, is activated in DRG neurons in diabetic neuropathy (Inceoglu et al., 2015), an effect that is likely mediated by the toxic end glycation byproduct methylglyoxal (Barragan-Iglesias et al., 2019). We initially hypothesized that type I IFNs might induce eIF2α phosphorylation in DRG neurons via PKR activation given the well-established induction of this pathway in other cell types (Pindel and Sadler, 2011). We did not find evidence for type I IFN-induced PKR-eIF2α signaling in DRG neurons, even over long time courses. Instead, we observed clear evidence for rapid activation of MNK1-eIF4E signaling in DRG neurons *in vitro* and *in vivo*. Signaling via eIF4E was also critical for the production of mechanical pain responses by type I IFNs and poly (I:C), which produces endogenous type I IFN production (Yamamoto et al., 2003; Kawai and Akira, 2008). Activation of MNK-eIF4E signaling by type I IFNs has been observed in other cell types where it has been linked to increased immune surveillance (Joshi et al., 2009). Multiple previous studies have demonstrated that activation of MNK-eIF4E-mediated translation events are causative in the production of chronic pain states, including neuropathic pain (Moy et al., 2017; Megat et al., 2019; Shiers et al., 2019). Since both viral infections and prolonged production of type I IFNs can cause neuropathic pain, it is possible that type I IFN receptor signaling to MNK-eIF4E may be a key pathway for production of these types of neuropathies.

## ACKNOWLEDGMENTS

We would like to thank Dr. Michael Iadarola (NIH) for advice on the experimental strategy using the synthetic dsRNA, poly (I:C). This work was supported by NINDS/NIH NS065926 (TJP), R01NS100788 (ZTC), and CONACyT doctoral fellowship program (UF-E). Cartoons in figure 6 were partially drawn using biorender.com.

## AUTHOR CONTRIBUTIONS

PB-I, UF-E, VJ and AW performed the experiments and analyzed data. TJP and PB-I conceived the original idea. TJP, PB-I, GD, VG-S and ZTC supervised the experiments and wrote the manuscript with the contribution of all coauthors.

## DECLARATION OF INTERESTS

The authors declare no competing interests.

